# Type I-F CRISPR-Cas distribution and array dynamics in *Legionella pneumophila*

**DOI:** 10.1101/276188

**Authors:** Shayna R. Deecker, Alexander W. Ensminger

**Affiliations:** Department of Biochemistry, University of Toronto, Toronto, Ontario, Canada; Department of Molecular Genetics, University of Toronto, Toronto, Ontario, Canada

## Abstract

In bacteria and archaea, several distinct types of CRISPR-Cas systems provide adaptive immunity through broadly similar mechanisms: short nucleic acid sequences derived from foreign DNA, known as spacers, engage in complementary base pairing with invasive genetic elements setting the stage for nucleases to degrade the target DNA. A hallmark of type I CRISPR-Cas systems is their ability to acquire spacers in response to both new and previously encountered invaders (naïve and primed acquisition, respectively). Our phylogenetic analyses of 47 *L. pneumophila* type I-F CRISPR-Cas systems and their resident genomes suggest that many of these systems have been horizontally acquired. These systems are frequently encoded on plasmids and can co-occur with nearly identical chromosomal loci. We show that two such co-occurring systems are highly protective and undergo efficient primed acquisition in the lab. Furthermore, we observe that targeting by one system’s array can prime spacer acquisition in the other. Lastly, we provide experimental and genomic evidence for a model in which primed acquisition can efficiently replenish a depleted type I CRISPR array following a mass spacer deletion event.

## Introduction

Microorganisms have evolved over millions of years to survive in harsh environments, backed in part by immune strategies that protect against foreign genetic elements, such as viral phages and foreign DNA elements (van Houte *et al.* 2016). Clustered regularly interspaced short palindromic repeats (CRISPR) coupled with associated *cas* genes form a potent adaptive immune response in numerous prokaryotic species (Barrangou *et al.* 2007; Brouns *et al.* 2008; Marraffini and Sontheimer 2008). These systems have been classified into six major types, which are further divided into various sub-types, based on their mechanism of action and Cas protein content (Makarova *et al.* 2011; 2015; Koonin *et al.* 2017).

A CRISPR-Cas response to invading DNA occurs in three distinct phases: adaptation, expression and interference (Barrangou *et al.* 2007; Brouns *et al.* 2008; Marraffini and Sontheimer 2008). In the adaptation phase, the CRISPR-Cas system acquires a DNA sequence (spacer) from the invader and integrates it into an array of spacers interspersed with repetitive sequences (Ishino *et al.* 1987; Mojica *et al.* 2000; Jansen, van Embden, Gaastra, and Schouls 2002b; Barrangou *et al.* 2007; Jackson *et al.* 2017). The spacers are generally derived from foreign elements whose infection was unsuccessful, such as defunct phage (Hynes *et al.* 2014), and form the basis of immunological memory for the bacterium. During the expression phase, the array is transcribed and processed to form CRISPR RNA (crRNA) molecules which form a surveillance complex with Cas proteins (Brouns *et al.* 2008; Haurwitz *et al.* 2010). Infection by an invading genetic element initiates the interference step, wherein the surveillance complex recognizes and binds the foreign DNA via base-pairing with the complementary crRNA and cleaves it, effectively neutralizing the threat to the host (Brouns *et al.* 2008; Jore *et al.* 2011; Wiedenheft *et al.* 2011; Westra *et al.* 2012).

Despite the sophistication of CRISPR-Cas systems, phages and other foreign DNA elements can still escape CRISPR-Cas targeting (Samson *et al.* 2013; Stanley and Maxwell 2018). A common mechanism of escape is the accumulation of random mutations within the foreign element which prevent complementary base pairing with crRNAs during interference (Deveau *et al.* 2008; Semenova *et al.* 2011; Datsenko *et al.* 2012). CRISPR-Cas systems can overcome this escape by acquiring new spacers; in fact, imperfect CRISPR targeting often leads to a highly efficient “primed” acquisition response, providing an intrinsic mechanism to protect against mutational escape (Swarts *et al.* 2012; Datsenko *et al.* 2012; Savitskaya *et al.* 2013; Fineran *et al.* 2014; Richter *et al.* 2014). Primed acquisition has been studied in type I-B (Li, Wang, Zhao, *et al.* 2014; Li, Wang, and Xiang 2014; Li *et al.* 2017), I-C (Rao *et al.* 2017), I-E (Swarts *et al.* 2012; Datsenko *et al.* 2012; Savitskaya *et al.* 2013; Fineran *et al.* 2014; Xue *et al.* 2015; Jackson *et al.* 2019) and I-F (Richter *et al.* 2014; Vorontsova *et al.* 2015; Staals *et al.* 2016; Heussler *et al.* 2016; Jackson *et al.* 2019) CRISPR-Cas systems. Interference-driven acquisition, or targeted acquisition, has also been observed in type I-C (Rao *et al.* 2017) and I-F systems (Staals *et al.* 2016), wherein a primed acquisition response occurs against a target with a perfect match to a spacer already within the array.

*Legionella pneumophila* is a gram-negative bacterium and the causative agent of Legionnaires’ disease (Brenner *et al.* 1979). Most isolates possess any of three different CRISPR-Cas systems: type I-C, I-F and/or II-B (D’Auria *et al.* 2010; Rao *et al.* 2016). Our lab has recently shown that all three types of CRISPR-Cas systems found in *L. pneumophila* isolates are active (Rao *et al.* 2016) and we have characterized the targeted acquisition response for the type I-C system (Rao *et al.* 2017). While much of our work to date has focused on the I-C systems of *L. pneumophila*, the type I-F systems of this pathogen are highly protective, remarkably diverse with respect to spacer content, and are frequently found on plasmids – suggesting that they may be circulated via horizontal gene transfer (Rao *et al.* 2016). In this study, we perform the first comprehensive phylogenetic analysis of the *L. pneumophila* type I-F systems and test a model by which horizontal acquisition of a mobile type I-F CRISPR-Cas system could replenish a collapsed chromosomal array.

## Results

### Phylogenetic analyses suggest widespread horizontal exchange of *L. pneumophila* type I-F systems

In order to explore the hypothesis that plasmid-based type I-F CRISPR-Cas systems in *L. pneumophila* could be circulated via horizontal gene transfer, we bioinformatically examined the diversity of type I-F CRISPR-Cas systems within this species. Leveraging both CRISPRCasFinder (Couvin *et al.* 2018) and CRISPRDetect (Biswas *et al.* 2016), we surveyed 525 draft and 6 completed *L. pneumophila* genomes. In total, we identified 47 *L. pneumophila* isolates that possessed type I-F systems (Table S1), including 5 that we had described previously (Rao *et al.* 2016). We next performed two types of phylogenetic analysis: *cas1* phylogeny (Fig 1A), which placed each CRISPR-Cas system into one of six phylotypes; and core-genome phylogeny (Fig 1B), which reflects the overall relatedness between each of the 47 isolates. A comparison of the two trees indicates a clear phylogenetic incongruence suggesting that horizontal acquisition has impacted the distribution of type I-F CRISPR-Cas systems within the species.

**Figure 1.**
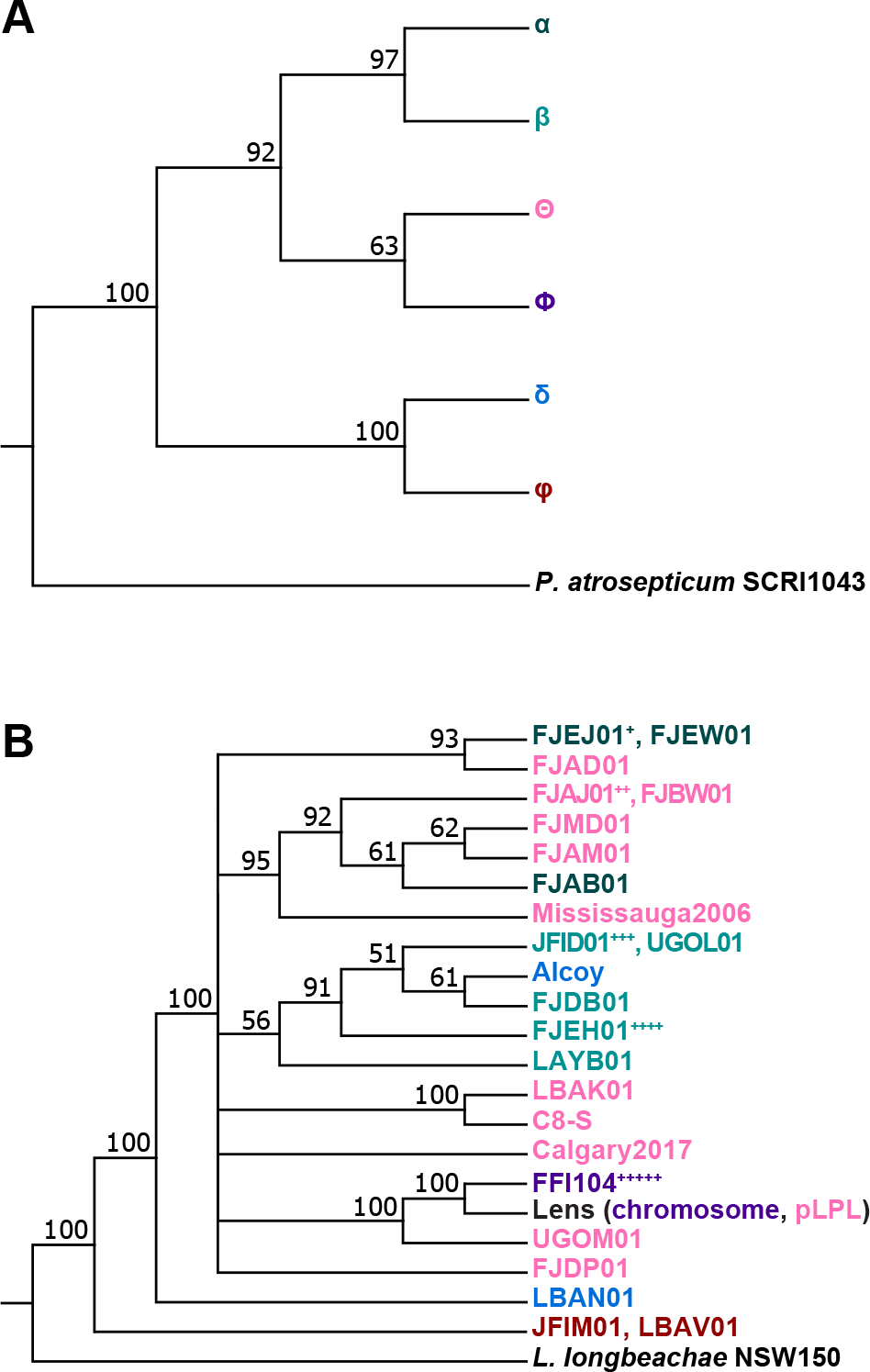
Phylogenetic analysis of type I-F CRISPR-Cas system diversity in *L. pneumophila* reveals a horizontal distribution across isolates. *L. pneumophila* draft and completed genomes were analyzed using CRISPRDetect and CRISPRCasFinder to identify type I-F systems present within the genomes. Isolates that possessed type I-F systems were subjected to phylogenetic analyses of their *cas1* gene and core genome. The core genome alignment for the examined isolates was determined by Roary. Isolate names are colour-coded based on the *cas1* gene phylogeny to allow for comparison between the analyses. The isolates used in this analysis and their accession numbers can be found in supplementary table 1. **A)** The *cas1* gene phylogeny for *L. pneumophila* isolates with type I-F systems reveals six different *cas1* groups. **B)** The core genome phylogeny of the examined isolates is not congruent with the *cas1* phylogeny, suggesting that the type I-F system was horizontally distributed rather than vertically inherited. Note that Lens possesses two type I-F CRISPR-Cas systems, one on a plasmid (group Θ) and one on its chromosome (group Φ). Also note that isolates with 100% nucleotide identity in their core genome, as well as a shared *cas* gene group and redundant CRISPR array, have been collapsed to one representative in the phylogeny. Isolates with unique CRISPR arrays are listed. Collapsed isolates include: +: FJEK01, FJEL01, FJEM01, FJEN01, FJEO01, FJEP01, FJEQ01, FJER01, FJES01, FJET01, FJEU01, FJEV01, FJEX01, FJEY01, FJEZ01, FJFA01 ++: FJBI01, FJBU01 +++: LAXR01 ++++: FJEI01 +++++: FFI105, FFI337

Given the results of these phylogenetic analyses, we next examined the spacer distribution across each of the arrays to determine the level of array diversification within each of the six *L. pneumophila* type I-F *cas1* phylotypes. We aligned, clustered, and visualized each of the 26 distinct *L. pneumophila* type I-F CRISPR arrays using CRISPRStudio (Dion *et al.* 2018) (Fig. 2). These analyses revealed patterns consistent with both spacer acquisition and spacer loss, suggesting that both processes contribute to *L. pneumophila* type I-F CRISPR array diversity.

**Figure 2.**
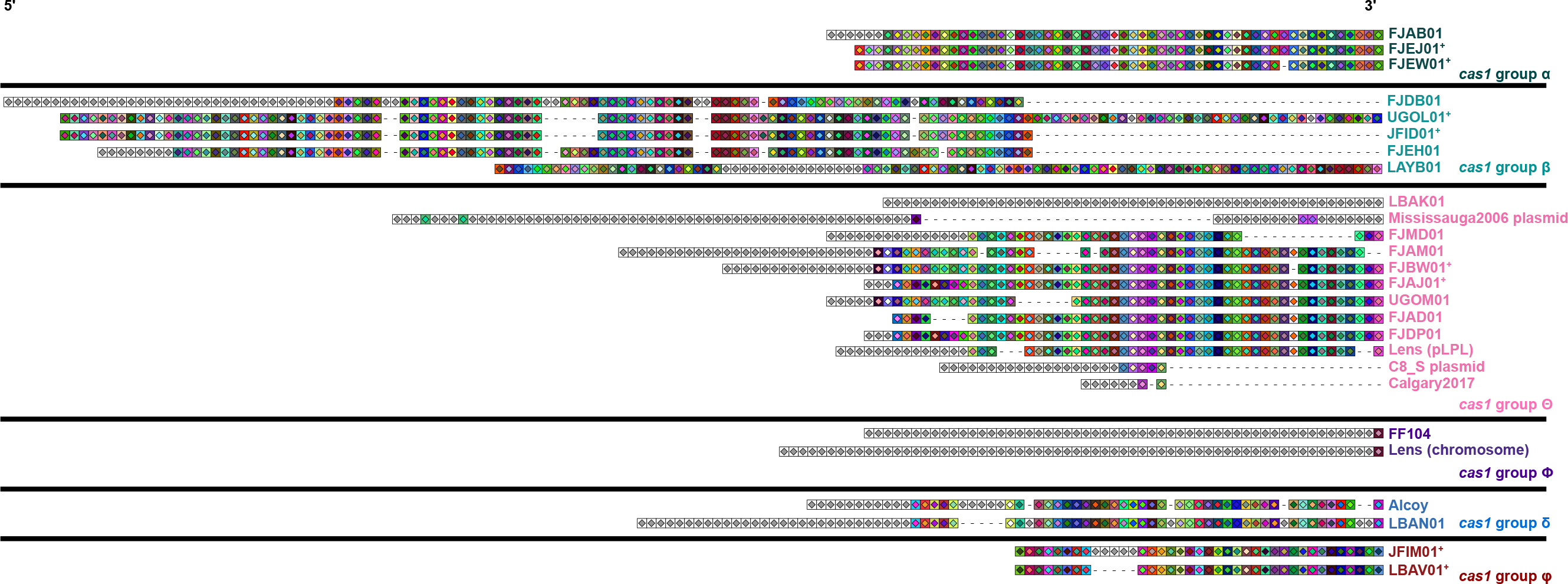
CRISPR array analysis suggests that spacer acquisition and spacer loss has contributed to array diversification in *L. pneumophila* type I-F systems. *L. pneumophila* isolates were subjected to a CRISPRStudio analysis to look at the spacer composition of their CRISPR arrays. Grey boxes denote unique spacers, coloured boxes denote shared spacers and a dashed line denotes spacer loss. The isolate colour coding scheme is based on the *cas1* grouping from figure 1. Denoted isolates within the same *cas* gene group have 100% nucleotide identity in their core genomes. Note that *cas1* group β appears to have spacer rearrangements present, resulting in an imperfect alignment between the arrays.

### The plasmid and chromosomal Lens CRISPR-Cas systems are active and adaptive

Given the evidence for spacer gain and loss that we observe in *L. pneumophila* type I-F arrays, (Fig. 2), we next decided to examine array dynamics experimentally. We focused on *L. pneumophila* str. Lens, which possesses two type I-F CRISPR-Cas systems: one on its chromosome and one on an endogenous 60 Kb plasmid, pLPL (D’Auria *et al.* 2010; Rao *et al.* 2016). The two systems have a 97.6% Cas protein identity and the repeat units between the spacers in the CRISPR array differ by only a single nucleotide (Rao *et al.* 2016) (Fig. 3). The CRISPR arrays themselves are of different lengths (64 spacers for the chromosomal system and 53 for that contained on the pLPL plasmid). Despite a high degree of identity between the Cas proteins, each array contains a completely unique set of spacers (D’Auria *et al.* 2010; Rao *et al.* 2016). The presence of two remarkably similar I-F systems in *L. pneumophila* str. Lens provided us with an opportunity to examine targeted spacer acquisition in both of these largely uncharacterized CRISPR-Cas systems and the interplay between them.

**Figure 3.**
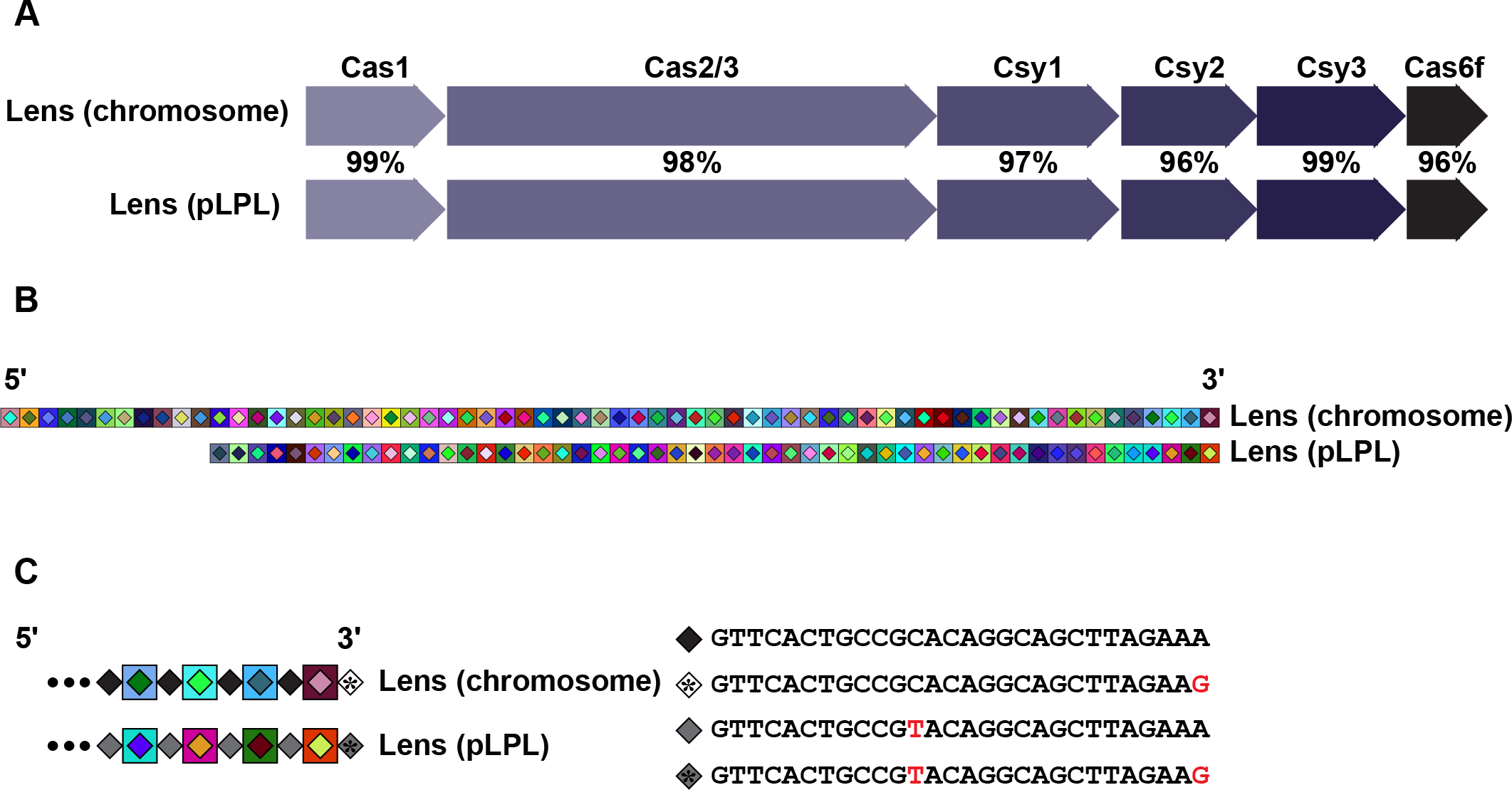
A comparison of the Lens chromosome and pLPL CRISPR-Cas systems. **A)** The overall pairwise amino acid identity across the Cas proteins of the two systems is approximately 97%, with individual pairwise Cas protein identities ranging from 96% to 99%. **B)** A CRISPRStudio alignment of the arrays for two systems shows completely unique spacer content and a differing number of spacers. **C)** Analysis of the repeat sequences shows that the Lens pLPL and chromosome systems have one SNP between their consensus repeats, in addition to possessing a mutated last repeat in their arrays. Mutations are denoted in red.

To assess CRISPR-Cas activity in both Lens type I-F systems, we performed an established transformation efficiency assay (Marraffini and Sontheimer 2008) using two different targeted protospacer sequences: one matching the most recently acquired spacer and one matching a spacer from the middle of the array (chromosomal spacer 23 and pLPL spacer 50). Consistent with active CRISPR-Cas protection, each of the protospacer-containing plasmids exhibited significant reductions in transformation efficiencies relative to a scrambled protospacer control (Fig. S1). These relative transformation efficiencies ranged from 1×10^−2^ to 1×10^−4^, with the most recently acquired spacers providing ~100-fold greater protection than spacers located in the middle of each array.

To determine whether spacer acquisition occurs within the context of a perfectly matched protospacer target, we pooled the transformed populations, passaged them on an automated liquid handler for 20 generations without selection, extracted their genomic DNA, and screened the leader end of the CRISPR array by PCR and agarose gel electrophoresis. While the populations transformed with plasmids encoding either protospacer 23 (chromosome) or protospacer 50 (pLPL plasmid) exhibited spacer acquisition in both Lens systems (Fig. 4), the populations transformed with protospacer 1 plasmids exhibited spacer loss, with spacer acquisition undetectable on a gel (Fig. 4). These data support our bioinformatic analyses, which indicated that both spacer acquisition and spacer loss contribute to type I-F CRISPR array diversity in *L. pneumophila* isolates (Fig. 2).

**Figure 4.**
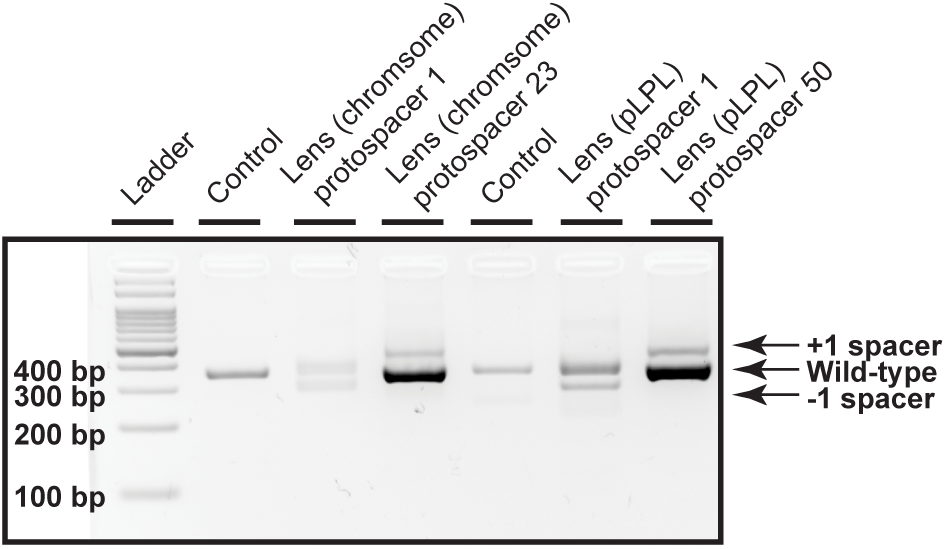
Spacer acquisition and spacer loss in both Lens CRISPR-Cas systems. Spacer acquisition and loss were analyzed using a PCR-based screen where the leader-end of the CRISPR array for both the control samples and the transformed samples was amplified with system-specific primers to differentiate between the chromosomal Lens and the plasmid Lens arrays and visualized on an agarose gel. Products from the transformed samples were compared to the control, which contained untransformed genomic DNA. Bands representing spacer acquisition and loss are indicated.

Our observation that the mid-array spacers of each type I-F system provided relatively modest protection led us to ask whether these spacers could nevertheless drive primed acquisition of new, more protective spacer sequences. To characterize the patterns of targeted spacer acquisition in the chromosomal and pLPL CRISPR-Cas systems, we amplified the leader-proximal region of each CRISPR array from wild-type populations that had been transformed with the plasmids targeted by their relatively permissive mid-array spacers (chromosomal: spacer 23; pLPL: spacer 50). We Illumina sequenced these PCR products and used an established bioinformatics pipeline (Rao *et al.* 2017) to identify newly acquired spacer sequences within each read (Table 1). We then mapped the target of each new spacer to the priming plasmid (Krzywinski *et al.* 2009) (Fig. 5).

**TABLE 1.**
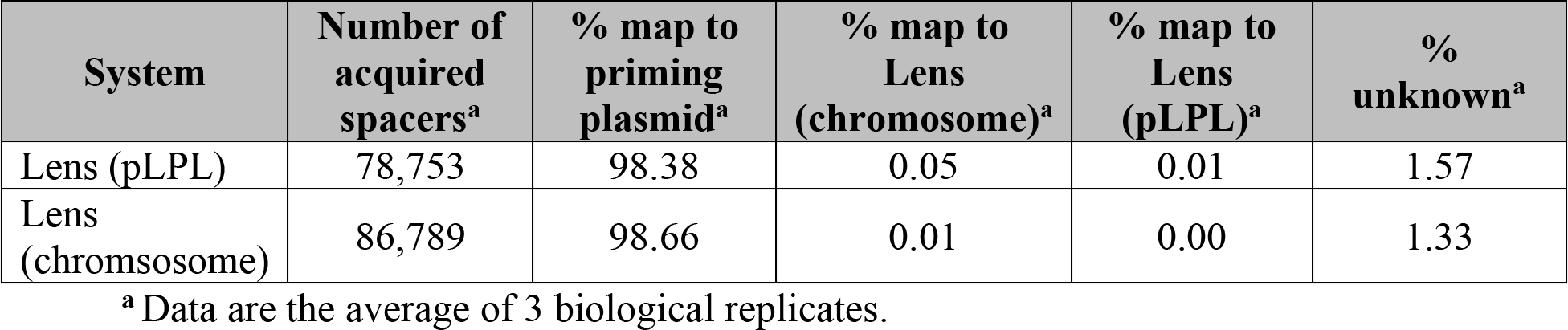
The number of acquired spacers for the Lens (pLPL) and Lens (chromosome) CRISPR-Cas systems and their respective targets during self-priming.

| **System** | **Number of acquired spacers<sup>a</sup>** | **% map to priming plasmid<sup>a</sup>** | **% map to Lens (chromosome)<sup>a</sup>** | **% map to Lens (pLPL)<sup>a</sup>** | **% unknown<sup>a</sup>** |
| --- | --- | --- | --- | --- | --- |
| Lens (pLPL) | 78,753 | 98.38 | 0.05 | 0.01 | 1.57 |
| Lens (chromosome) | 86,789 | 98.66 | 0.01 | 0.00 | 1.33 |

<sup>a</sup> Data are the average of 3 biological replicates.

**Figure 5.**
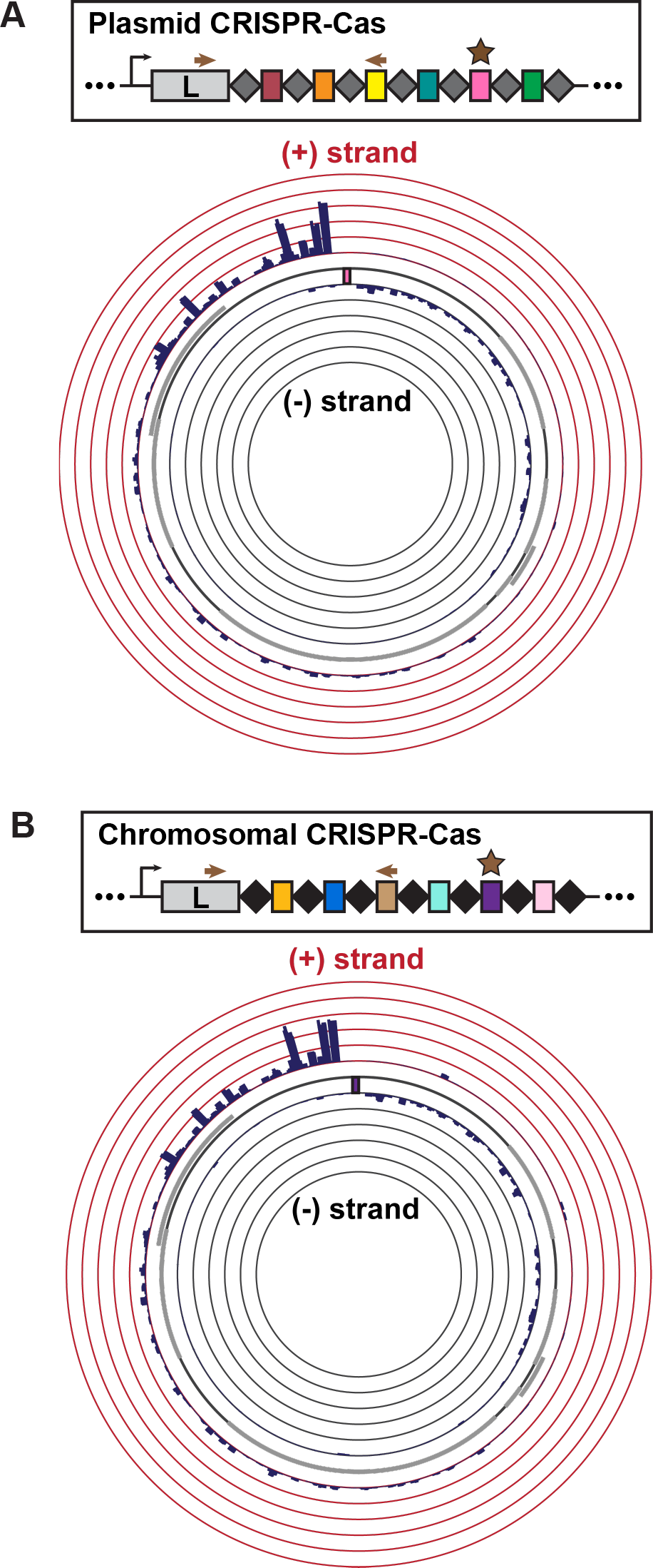
Characterization of self-primed spacer acquisition in the two Lens CRISPR-Cas systems. Bacterial transformants with targeted plasmids were passaged for 20 generations without antibiotic selection to enrich for spacer acquisition; the leader end of the CRISPR array was amplified and the amplicons were Illumina sequenced. Newly targeted protospacers were obtained from the raw reads using an in-house bioinformatics pipeline and visualized with Circos. The arrows in the simplified array schematic show the primer location for PCR amplification prior to sequencing, while the star denotes the priming spacer. All data are the average of three biological replicates. **A)** The distribution of newly targeted protospacers mapped to the priming plasmid on the Circos plot reveals a strand bias in self-primed spacer acquisition within the Lens (pLPL) CRISPR-Cas system. The height of the bars indicates the number of spacers mapped to the position on the plasmid, up to 5% of total acquired spacers. **B)** The distribution of newly targeted protospacers mapped to the priming plasmid shows a similar pattern of self-primed spacer acquisition within the Lens (chromosome) CRISPR-Cas system to that of its pLPL CRISPR-Cas counterpart. Labelling as in **(A)**.

Both the Lens type I-F CRISPR-Cas systems exhibited a biased distribution of acquired spacers (Fig. 5, S2) consistent with what has been seen previously for the type I-F systems of *Pectobacterium astrosepticum* (Richter *et al.* 2014; Staals *et al.* 2016) and *Pseudomonas aeruginosa* (Vorontsova *et al.* 2015; Heussler *et al.* 2016). The majority of the newly targeted protospacers clustered around the priming sequence on the targeted plasmid, with the non-primed strand of DNA (the plus (+) strand) containing ~75% of these protospacers (Fig. 5). Consistent with observations in other type I-F systems (Richter *et al.* 2014; Staals *et al.* 2016; Heussler *et al.* 2016), switching the target sequence to the opposite strand led to an acquisition pattern that mirrored the original distribution observed when the (-) strand contained the targeted protospacer (Fig. S3). The spacer length distribution and PAM usage were also consistent with previous observations of other type I-F systems (Mojica *et al.* 2009; Cady *et al.* 2012; Richter *et al.* 2014; Vorontsova *et al.* 2015; Staals *et al.* 2016) (Fig. S3B, C, S4, S5). Taken together, these data suggest that spacer acquisition is qualitatively similar between the chromosomal and pLPL CRISPR-Cas systems.

### Permissive targeting by one system can lead to primed acquisition in the other system

Since the chromosomal and pLPL CRISPR-Cas systems function in a very similar manner during targeted acquisition and share a high degree of homology within their Cas genes and repeat sequences, we speculated that priming of one system might lead to spacer acquisition in the other. Specifically, we tested whether introducing a protospacer-containing plasmid targeted by one CRISPR-Cas system would initiate a primed acquisition response in the second system. Indeed, this led to efficient spacer acquisition on the second array (Fig. 6), with patterns largely indistinguishable from what we previously observed on the cognate array (Fig. 5).

**Figure 6.**
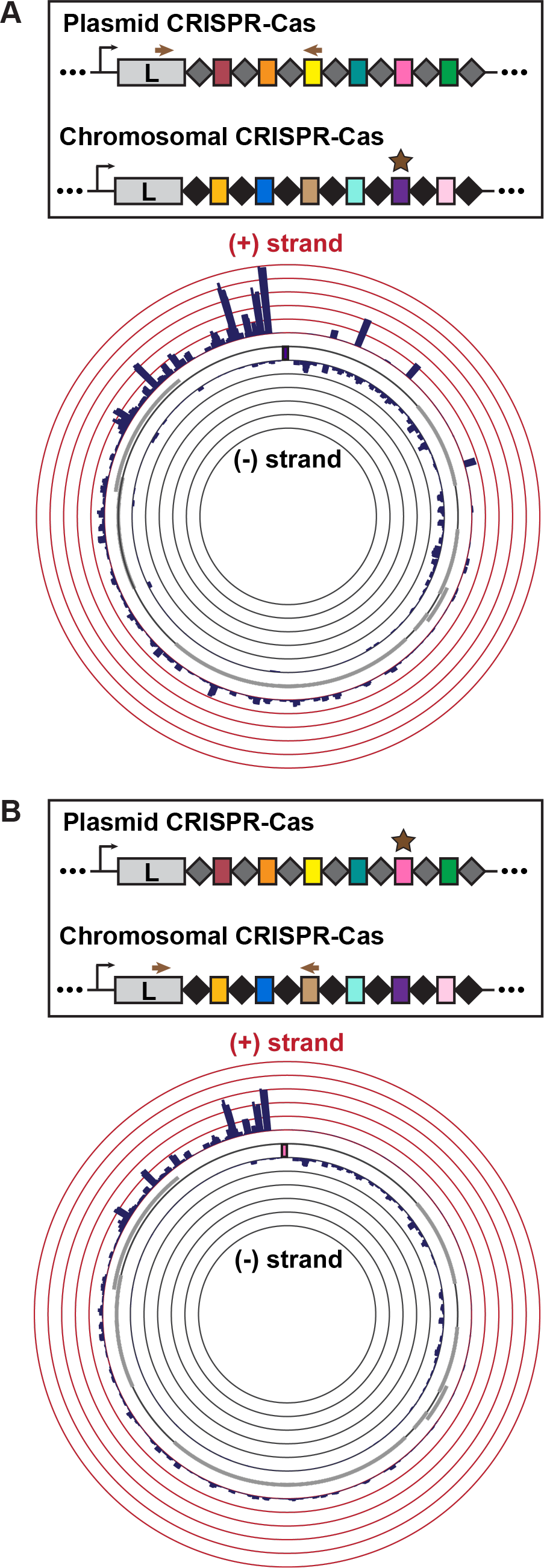
Characterization of cross-primed spacer acquisition in the two Lens CRISPR-Cas systems. The experimental set-up is the same as described in figure 5. All data are the average of three biological replicates. The distribution of newly targeted protospacers mapped to the priming plasmid reveals cross-priming between the Lens (chromosome) and Lens (pLPL) CRISPR-Cas systems. **(A)** shows Lens (chromosome) primed, Lens (pLPL) array examined while **(B)** shows Lens (pLPL) primed, Lens (chromosome) array examined.

### Primed repopulation of collapsed arrays

Based on our observations suggesting widespread horizontal inheritance of *L. pneumophila* type I-F systems (Fig. 1), the diversity of spacer sequences (Fig. 2), and the ability of closely related systems to prime each other (Fig. 6), we next asked whether coincident CRISPR-Cas might provide a mechanism for replenishing collapsed chromosomal arrays. Spacer loss is one of several outcomes when the targeting of a particular sequence becomes detrimental to bacterial survival. In the lab, this occurs when we artificially “force” the coexistence of an efficiently targeted plasmid and an active CRISPR-Cas system through selection (Fig. 4). Similar events are also likely to occur randomly or when CRISPR-Cas systems acquire self-targeting spacers at a low, but detectable rate (Fig. 7) (Yosef *et al.* 2012; Datsenko *et al.* 2012; Savitskaya *et al.* 2013; Vorontsova *et al.* 2015; Staals *et al.* 2016; Rao *et al.* 2017). In the most extreme instances, an entire array of spacers could be lost through recombination between the first and last repeats. Normally, such a loss could only be reversed by the relatively inefficient mechanism of naïve spacer acquisition (Yosef *et al.* 2012; Datsenko *et al.* 2012; Savitskaya *et al.* 2013). However, if such a collapsed array could be restored through primed acquisition driven by a coincident system, strains with multiple arrays would be inherently more resistant to the catastrophic loss of CRISPR-Cas protection. Such a model makes a number of predictions (Fig. 7), which we sought to bioinformatically and experimentally test.

**Figure 7.**
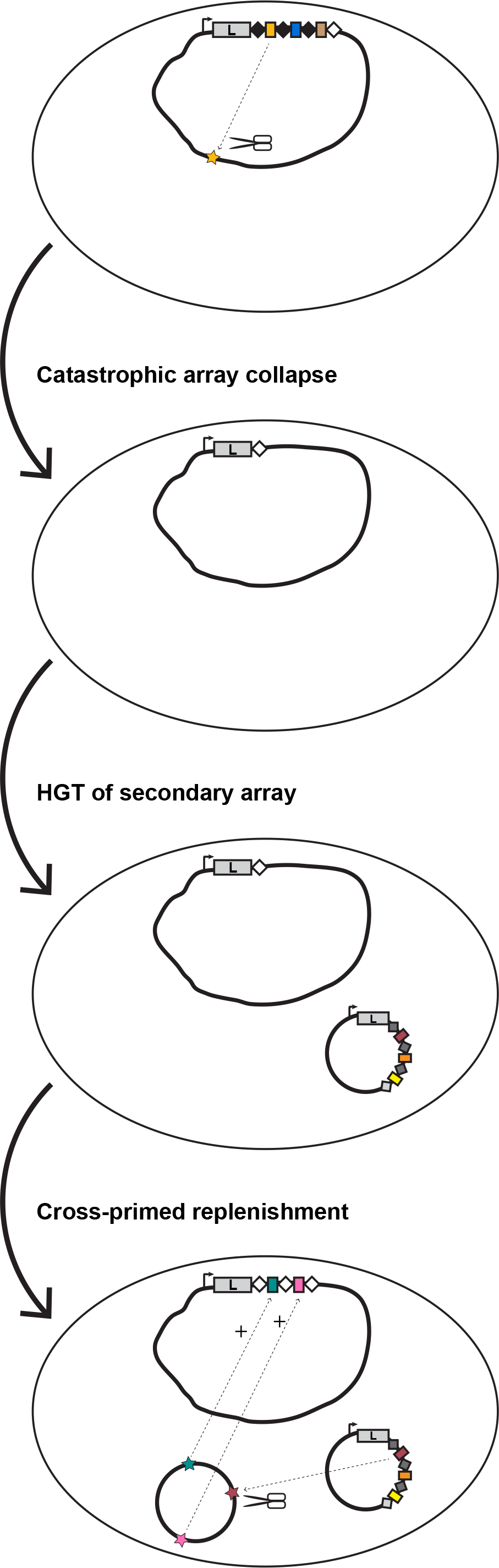
Cross-priming between two similar CRISPR-Cas systems can repopulate a collapsed CRISPR array. A bacterium with a CRISPR-Cas system could undergo a mass spacer deletion event through homologous recombination between the first (black) and last (white) repeat sequences. The remaining locus carries a single repeat (white). While such “catastrophic collapse” events are likely to occur randomly at a certain rate, a driver of such a collapse could be the acquisition of a self-targeting spacer (yellow), selecting for spacer loss. Horizontal acquisition of a second CRISPR-Cas array (e.g. on a plasmid) is a first step towards replenishing the primary array. If cross-priming can occur between this secondary array and the collapsed array, the original CRISPR array is replenished, but bears an observable molecular scar - conversion of all the repeats to the sequence of the last repeat (white).

First, we used allelic replacement to generate an *L. pneumophila* Lens strain in which the entire chromosomal array was replaced by a single copy of its last repeat, mimicking what would occur after complete spacer loss. Next, we transformed two independently derived array deletion strains with the pLPL type I-F priming plasmid as above. Using PCR and Illumina sequencing, we observed robust spacer acquisition in the formerly depleted chromosomal CRISPR array (Fig 8, Table S4), indicating that primed acquisition can replenish the completely collapsed array of a coincidental type I-F system. As expected, the consensus repeat sequence of this replenished array adopted the same alternate sequence as the last repeat of the array.

**Figure 8.**
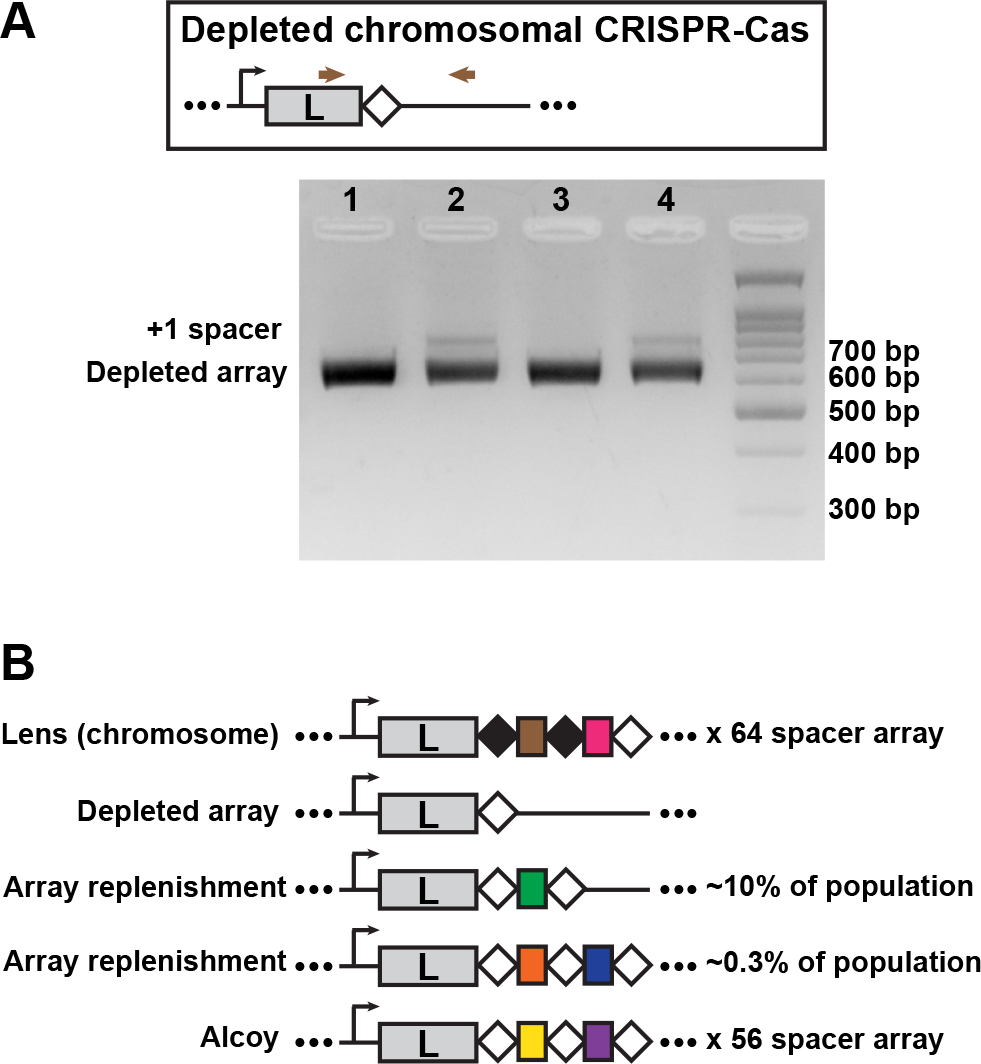
An experimentally depleted chromosomal CRISPR array can be replenished through the activity of a plasmid-based array. **A)** Replenishment of a depleted array in the lab. Allelic replacement was used to remove the entire array from the chromosomal Lens I-F system, leaving behind a single, last repeat sequence (see materials and methods). This strain was then transformed with a plasmid targeted by the pLPL (Lens plasmid-based) I-F array previously shown to drive primed acquisition (pLPL protospacer 50; see Figure 4). Spacer acquisition by the empty chromosomal array was analyzed using a PCR based screen where the leader-end of the CRISPR array was amplified and visualized on an agarose gel. Products from the transformed samples (samples 2 and 4) were compared to untransformed controls (samples 1 and 3). Samples 1 and 2 are from depleted array clone #1 and samples 3 and 4 are from depleted array clone #2. **B)** Repeat signatures of depletion/replenishment. The repeat structure of the experimentally replenished Lens CRISPR arrays resembles that of *L. pneumophila* str. Alcoy, suggesting a similar array depletion/replenishment event may have occurred within the Alcoy lineage. The frequency of acquiring one new spacer versus two new spacers following replenishment in the Lens array depletion isolates was determined by Illumina sequencing.

We next examined the CRISPR (repeat) sequences in each of the isolates used in our earlier bioinformatic analyses for similar evidence of complete collapse followed by replenishment. In many of the arrays, there is a consensus repeat that is found throughout the majority of the array, with the last repeat in the array carrying a mutation (Table 2). This has been observed before in other systems (Jansen, van Embden, Gaastra, and Schouls 2002a; 2002b; Horvath *et al.* 2008; Lopez-Sanchez *et al.* 2012). However, for a number of the *L. pneumophila* isolates, including Alcoy, JFIM01, LBAN01, and LBAV01, this is not the case. In these isolates, their consensus repeat (found throughout the CRISPR array) is identical to their last repeat. Intriguingly, the sequence that remains is identical to the last repeat found in other type I-F isolates – what one would predict if they were the product of complete array collapse followed by subsequent replenishment (Fig. 8).

**TABLE 2.**
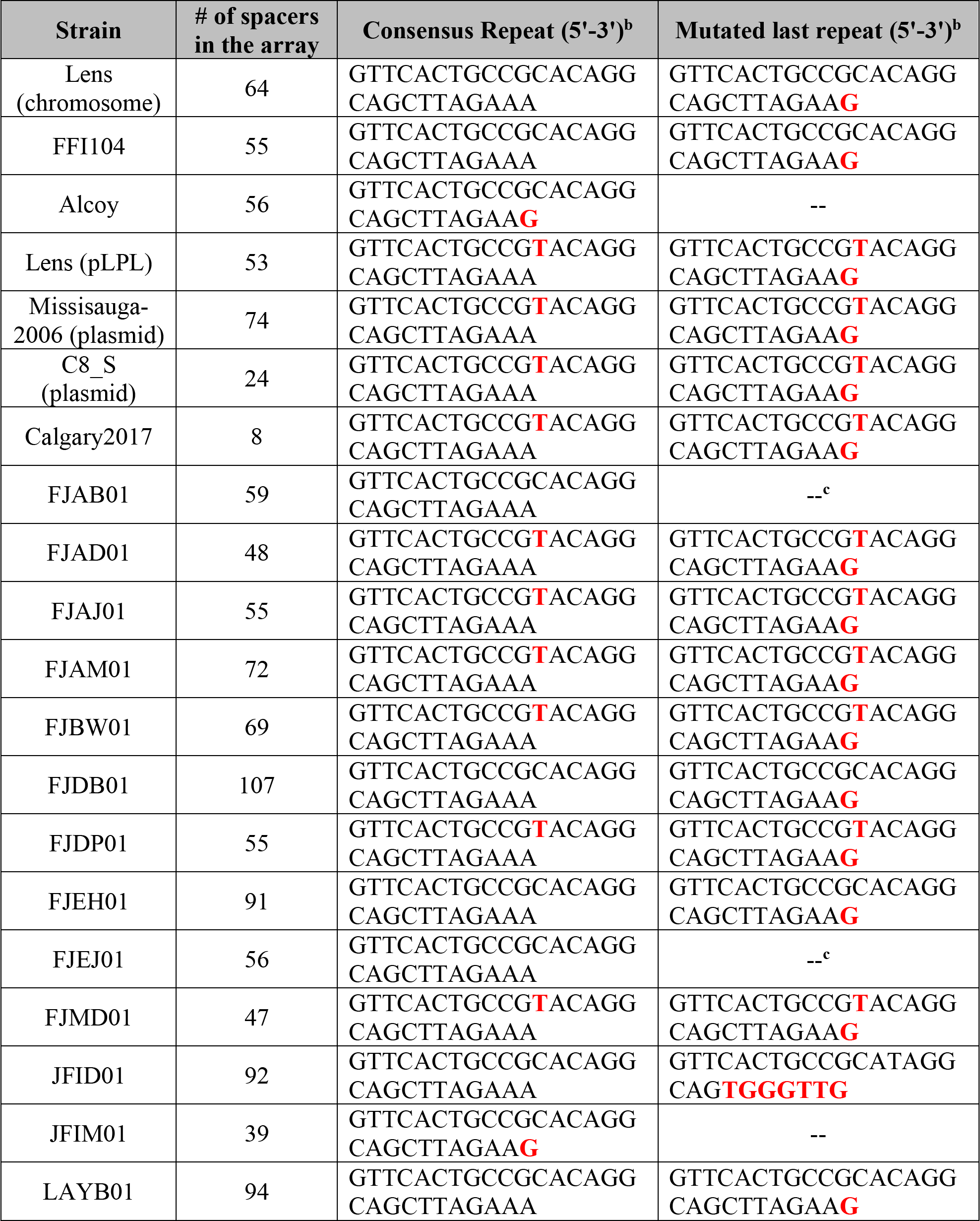
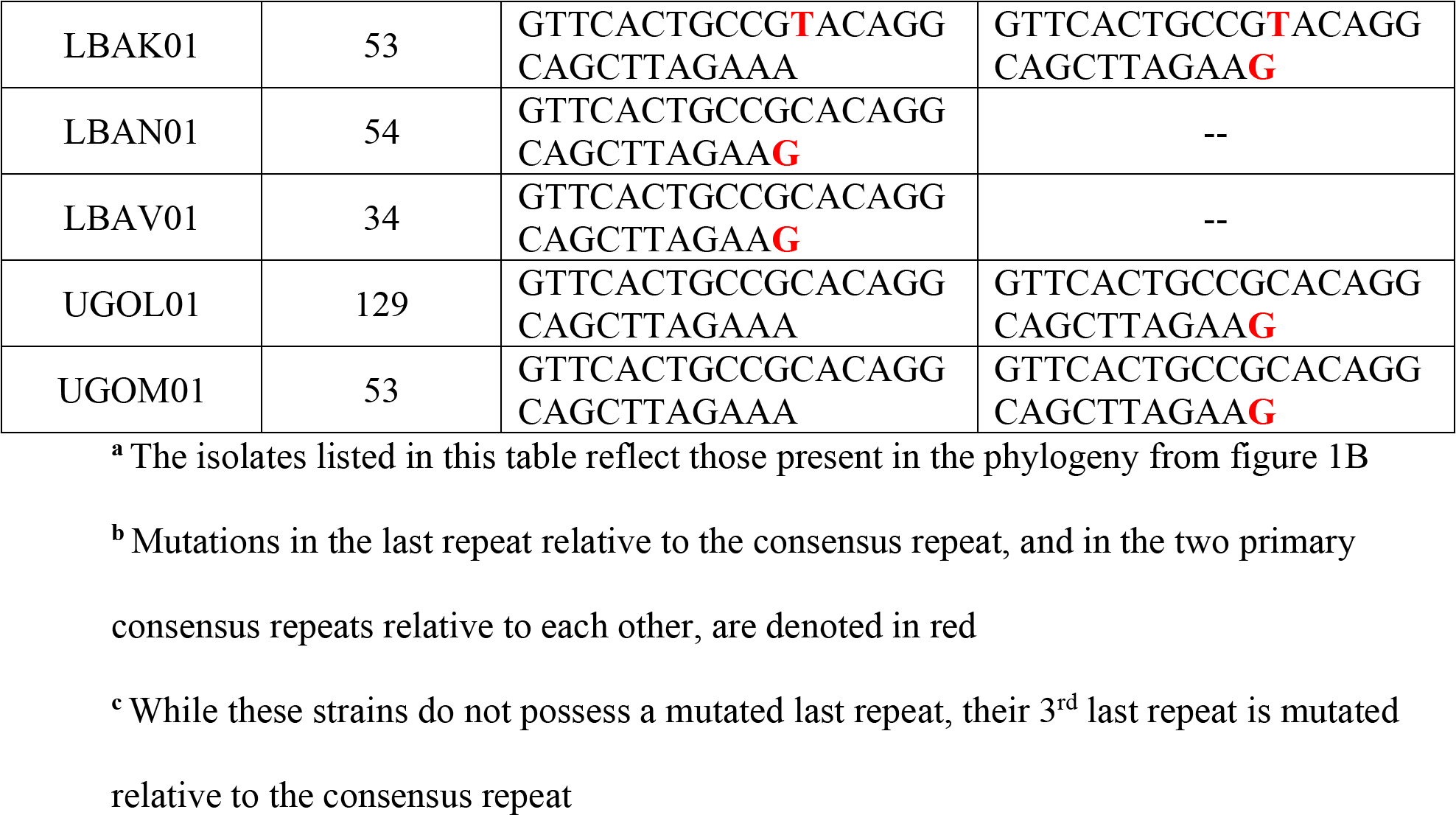
The repeat sequences of *L. pneumophila* type I-F CRISPR-Cas systems.^**a**^

## Discussion

Horizontal gene transfer is a driving force in shaping bacterial biology and pathogenicity (Ochman *et al.* 2000; Juhas 2013; Hall *et al.* 2017). With respect to *L. pneumophila*, comparative genomic studies have highlighted the importance of horizontal gene transfer, mobile genetic elements, and homologous recombination in shaping the bacterium’s evolutionary trajectory (Cazalet *et al.* 2004; 2008; D’Auria *et al.* 2010; Gomez-Valero *et al.* 2011; Sánchez-Busó *et al.* 2014; Gomez-Valero *et al.* 2014; McAdam *et al.* 2014; Burstein *et al.* 2016; David *et al.* 2017). This aspect of *L. pneumophila* biology extends to the presence and maintenance of plasmid-based CRISPR-Cas systems (D’Auria *et al.* 2010; Rao *et al.* 2016). Our bioinformatic analyses suggest that CRISPR-Cas type I-F systems are horizontally distributed in this species (Fig. 1). These CRISPR arrays have also undergone extensive spacer acquisition and some spacer loss (Fig. 2). Horizontal acquisition of type I-F systems is well-established in other species. For instance, in *Vibrio* species, 97% of identified type I-F systems are encoded on mobile genetic elements (McDonald *et al.* 2019). Like *L. pneumophila*, it’s hypothesized these systems have been acquired by their host through horizontal gene transfer (McDonald *et al.* 2019). Our data, however, suggest that such transfer is not merely a mechanism by which isolates acquire CRISPR-Cas protection, but also a potential mechanism to maintain existing defensive capabilities through the unique spacer dynamics provided by two inter-priming arrays. These inter-priming arrays can be part of two different systems, as demonstrated by our data, but could also occur in a system where two (or more) different arrays share a set of *cas* genes (Swarts *et al.* 2012; Datsenko *et al.* 2012; Staals *et al.* 2013; 2014; Majumdar *et al.* 2015; Elmore *et al.* 2015; Silas *et al.* 2017). For example, in *E. coli* str. K12, it has been shown that two different CRISPR arrays can be populated by the same set of *cas* genes (Swarts *et al.* 2012; Datsenko *et al.* 2012).

We propose that when a bacterium acquires a second, closely related CRISPR-Cas system, it gains a mechanism by which depleted CRISPR arrays can be repopulated (Fig. 7). We have modeled such an event in the lab, made predictions about what the signatures of such events would be, and provide genomic data to suggest it may have occurred on several occasions within our collection of sequenced isolates. One obvious line of future investigation would be to observe whether such patterns are present in other species with mobilized type I CRISPR-Cas and perhaps absent in instances where CRISPR-Cas is acquired primarily through vertical inheritance.

Lastly, while we think of spacer loss as a predominantly negative event (loss of protection), it is likely to play a more nuanced role in the maintenance of CRISPR-Cas activity. Clearly, array length is the product of a dynamic process whose impact on adaptation, expression, and interference remains largely unexplored. Many of the type I-F systems in *L. pneumophila* have different array lengths, ranging from 8 spacers to 129 spacers, with an average length of 61 spacers (Table 2). Toms and Barrangou have performed a global analysis of class I CRISPR arrays and found that the average array length for type I-F systems was 33 spacers, with statistically significant differences between the array lengths of different type I subtypes (Toms and Barrangou 2017). Accordingly, if spacer acquisition is a driving force in array divergence, it is likely coupled to spacer loss. Close examination of the mechanisms driving spacer loss in these systems - and their subsequent impact on CRISPR-Cas functionality - will be crucial to further testing the model of array diversification in *L. pneumophila*.

## Methods and Materials

### Bioinformatic analyses

Bioinformatic analyses of the Illumina sequence data were performed as described previously (Rao *et al.* 2017). Briefly, the raw paired-end reads were merged using FLASH (Magoc and Salzberg 2011), and any unpaired reads were subsequently quality trimmed using Trimmomatic (Bolger *et al.* 2014). These processed reads were then combined and analyzed using a Perl script (available upon request) that annotated existing spacers (S), newly acquired spacers (X), repetitive sequences (R) and the downstream sequence (D). The newly acquired spacers were aligned to the priming plasmid, the *L. pneumophila* str. Lens chromosome or the *L. pneumophila* str. Lens plasmid using BLASTN (Altschul *et al.* 1990). The results from the BLASTN alignment for the priming plasmid were then processed to obtain coverage per nucleotide, and plotted on the reference sequence using Circos (Krzywinski *et al.* 2009).

For the bioinformatic analyses of the *L. pneumophila* type I-F system diversity, *L. pneumophila* draft genomes and completed genomes were downloaded from the European Nucleotide Archive and NCBI respectively. Type I-F CRISPR-Cas systems were identified using CRISPRCasFinder (Couvin *et al.* 2018) and CRISPRDetect (Biswas *et al.* 2016). All genomes with type I-F systems present were annotated using Prokka (Seemann 2014). For the core genome phylogeny, pan-genome analysis was performed with Roary(Page *et al.* 2015). Both the *cas1* and core genome phylogenies were created with RAxML (Stamatakis 2014) and the RAxML trees were condensed with MEGA7 (Kumar *et al.* 2016). CRISPR array alignments, clustering and visualization were performed with CRISPRStudio (Dion *et al.* 2018).

### Bacterial strains, plasmids and oligos used

The bacterial strains and plasmids used in this study are listed in supplementary table 2, and the oligos used in this study are listed in supplementary table 3.

The priming plasmids were created by annealing oligos (see Table S3) to create the protospacer insert with the canonical GG PAM (Mojica *et al.* 2009; Cady *et al.* 2012; Richter *et al.* 2014; Vorontsova *et al.* 2015; Staals *et al.* 2016) and subsequently ligating the insert into an ApaI/PstI-digested pMMB207 vector (Solomon *et al.* 2000). The scrambled control plasmid was created in the same manner, except it contained a 32-nt scrambled sequence in place of a targeted protospacer sequence.

Lens (chromsome) array deletion mutants were generated through allelic replacement. Briefly, 1 kb of DNA upstream and 2 kb downstream of the CRISPR array were amplified by PCR and stitched together to create an insert where the entire array, save for the last repeat, was deleted. The insert was ligated into a pJB4648 plasmid. Overnight cultures of *L. pneumophila* str. Lens were grown in ACES-buffered yeast extract (AYE) medium to an OD_600_ of ~4.0 using two-day patches that were grown on charcoal-buffered ACES yeast extract (CYE) plates. Pellets from 4.0 ODU of culture underwent three washing steps: twice with 1 mL of ice-cold ultrapure water and once with 1 mL of ice-cold 10% glycerol. The pellet was then re-suspended in 200 uL of ice-cold 10% glycerol and 400 ng of plasmid was added to the sample. The solution was transferred to an ice-cold electroporation cuvette with a 2 mm gap and electroporated with the following settings: 2500 kV, 600 Ω and 25 mF. After electroporation, 800 uL of AYE medium was added to each sample and the samples recovered for 3 hours at 37°C at 600 RPM in a shaking incubator. The samples were plated on CYE plates supplemented with 15 μg mL^−1^ of gentamycin and incubated at 37°C for 3 days. Surviving colonies were patched onto CYE + gentamycin plates and grown at 37°C for 2 days. Patches were subsequently struck onto CYE plates supplemented with sucrose and incubated at 37°C for 3 days. Surviving colonies were patched onto CYE + sucrose plates, grown at 37°C for 2 days and screened by PCR to confirm the deletion. Two independent clones were Illumina sequenced and used for subsequent replenishment assays.

### Transformation efficiency assay and population pool generation

The transformation efficiency assay was performed as we have described previously (Rao *et al.* 2016) with some modifications. Briefly, *L. pneumophila* str. Lens was electroporated as described above. The samples were plated in a dilution series on CYE plates supplemented with 5 μg mL^−1^ of chloramphenicol and incubated at 37°C for 3 days. The relative transformation efficiency for each targeted plasmid was calculated as a percentage of the transformation efficiency obtained from the scrambled control plasmid. Three biological replicates were performed for each transformation efficiency assay.

Population pools for spacer acquisition experiments were generated by mixing together ≥ 50 colonies per population from a newly transformed wild-type strain on CYE plates supplemented with 5 μg mL^−1^ of chloramphenicol using AYE medium supplemented with 5 μg mL^−1^ of chloramphenicol. Population pools were made in triplicate for each transformed plasmid.

### Serial passaging on an automated liquid handler

The serial passaging of transformed *L. pneumophila* str. Lens populations was performed as described previously (Rao *et al.* 2016). Briefly, overnight cultures of the population pools in AYE medium supplemented with 5 μg mL^−1^ of chloramphenicol for plasmid maintenance were grown to an OD_600_ of ~2.0. The culture was then back diluted to an OD_600_ of ~0.0625 and grown in a flat-bottom 48-well plate (Greiner) in a shaking incubator at 37°C. A Freedom Evo 100 liquid handler (Tecan) connected to an Infinite M200 Pro plate reader (Tecan) measured the optical density of the plate every 20 minutes, until an OD_600_ of ~2.0 was reached. The cultures were then automatically back diluted to an OD_600_ of ~0.0625 in the adjacent well to continue growth, and the remaining culture was transferred to a 48-well plate that was kept at 4°C. In this manner, each saved culture represented ~5 generations of growth. The passaging was done without selection in AYE medium to allow for plasmid loss during passaging.

### Genomic DNA extraction, PCR and agarose gel screening

Genomic DNA was extracted from the passaged cultures and the parental chromosome Lens array deletion strains using a Machery-Nagel Nucleospin Tissue kit according to the manufacturer’s protocol. The extracted samples from passaged cultures were used as a template in a 30-cycle PCR reaction with Econotaq Polymerase (Lucigen) to amplify the leader end of the CRISPR array using primers listed in Table S3. The PCR products were then separated on a 3% agarose gel to determine if spacer acquisition (or spacer loss) had occurred based on the presence of an upper (or lower) band relative to the control sample.

### Nextera library prep and Illumina sequencing

The extracted genomic DNA from passaged cultures was prepared for leader-end array sequencing by performing a 20-cycle PCR using Kapa HiFi Polymerase (Kapa Biosystems) and the primers listed in Table S3. The PCR products were purified using a Machery-Nagel Nucleospin Gel and PCR Clean-up kit as per the manufacturer’s instructions and normalized to 1 ng using the Invitrogen Quant-iT PicoGreen dsDNA assay. The DNA was then tagmented using a Nextera XT tagmentation kit as per the manufacturer’s instructions. The tagmented products were sequenced with a paired-end (2 × 150 bp) sequencing run on an Illumina NextSeq platform at the Centre for the Analysis of Genome Evolution and Function (CAGEF) at the University of Toronto.

The genomic DNA from the parental chromosome Lens array deletion strains was normalized to 1 ng using PicoGreen. The DNA was then tagmented using a Nextera XT tagmentation kit as per the manufacturer’s instructions. The tagmented products were sequenced with paired-end (2 × 150 bp) sequencing in-house on an Illumina MiniSeq platform.

## Data Accessibility

The raw Illumina reads have been deposited into the NCBI sequence read archive under the BioProject PRJNA433194.

## Acknowledgements

The authors thank Griffin Deecker (a volunteer high school student) for his assistance in bioinformatically examining the diversity of I-F repeat sequences in *L. pneumophila*, Chitong Rao for his contributions to experimental design and Kamran Rizzolo for discussions regarding phylogenetic analyses. We also thank the Center for the Analysis of Genome Evolution and Function (CAGEF) at the University of Toronto for performing Illumina sequencing. We thank members of the Ensminger laboratory for their suggestions and careful reading of the manuscript, in particular Beth Nicholson and Malene Urbanus. SRD is supported by a fellowship from the Department of Biochemistry, University of Toronto. This work was supported by a Project Grant from the Canadian Institutes of Health Research (PHT-148819), the Connaught Fund (NR-2015-16), and an infrastructure grant from the Canada Foundation for Innovation and the Ontario Research Fund (30364) to AWE.

## Notes

#### Summary of Updates

Changes to the manuscript include: 1) new, systematic phylogenetic analysis of L. pneumophila Type I-F CRISPR-Cas systems and their spacer content; 2) direct experimental evidence of primed replenishment following a completely collapsed array; 3) improved referencing and focus.

