## Supplemental Materials for "Type I-F CRISPR-Cas distribution and array dynamics in *Legionella pneumophila*"

### SUPPLEMENTARY TABLE 1

The isolates used in the bioinformatic analysis of type I-F CRISPR-Cas systems.

| Name | Accession Number | Database |
|---|---|---|
| *Legionella pneumophila* str. Alcoy | NC_014125.1 | Genbank |
| *Legionella pneumophila* str. C8_S | CP015939.1 (chromosome)<br>CP015940.1 (plasmid) | Genbank |
| *Legionella pneumophila* str. FFI104 | CP016872.1 | Genbank |
| *Legionella pneumophila* str. FFI105 | CP016873.1 | Genbank |
| *Legionella pneumophila* str. FFI337 | CP016876.1 | Genbank |
| *Legionella pneumophila* str. Lens | NC_006369.1 (chromosome)<br>NC_006366.1 (plasmid) | Genbank |
| *Legionella pneumophila* str. Calgary2017 | -- | Not yet deposited in a database |
| *Legionella pneumophila* str. Mississauga2006 | SRX1091506 | Sequence Read Archive |
| *Legionella pneumophila* str. 2531STDY5467307 | FJAB01 | European Nucleotide Archive |
| *Legionella pneumophila* str. 2531STDY5467313 | FJAD01 | European Nucleotide Archive |
| *Legionella pneumophila* str. 2531STDY5467293 | FJAJ01 | European Nucleotide Archive |
| *Legionella pneumophila* str. 2531STDY5467317 | FJAM01 | European Nucleotide Archive |
| *Legionella pneumophila* str. 2531STDY5467292 | FJBI01 | European Nucleotide Archive |
| *Legionella pneumophila* str. 2531STDY5467289 | FJBU01 | European Nucleotide Archive |
| *Legionella pneumophila* str. 2531STDY5467322 | FJBW01 | European Nucleotide Archive |
| *Legionella pneumophila* str. 2531STDY5467368 | FJDB01 | European Nucleotide Archive |
| *Legionella pneumophila* str. 2531STDY5467401 | FJDP01 | European Nucleotide Archive |
| *Legionella pneumophila* str. 2531STDY5467437 | FJEH01 | European Nucleotide Archive |
| *Legionella pneumophila* str. 2531STDY5467440 | FJEI01 | European Nucleotide Archive |
| *Legionella pneumophila* str. 2531STDY5467453 | FJEJ01 | European Nucleotide Archive |
| *Legionella pneumophila* str. 2531STDY5467426 | FJEK01 | European Nucleotide Archive |
| *Legionella pneumophila* str. 2531STDY5467455 | FJEL01 | European Nucleotide Archive |
| *Legionella pneumophila* str. 2531STDY5467459 | FJEM01 | European Nucleotide Archive |
| *Legionella pneumophila* str. 2531STDY5467458 | FJEN01 | European Nucleotide Archive |
| *Legionella pneumophila* str. 2531STDY5467461 | FJEO01 | European Nucleotide Archive |
| *Legionella pneumophila* str. 2531STDY5467462 | FJEP01 | European Nucleotide Archive |

| | | |
|---|---|---|
| *Legionella pneumophila* str. 2531STDY5467460 | FJEQ01 | European Nucleotide Archive |
| *Legionella pneumophila* str. 2531STDY5467463 | FJER01 | European Nucleotide Archive |
| *Legionella pneumophila* str. 2531STDY5467467 | FJES01 | European Nucleotide Archive |
| *Legionella pneumophila* str. 2531STDY5467468 | FJET01 | European Nucleotide Archive |
| *Legionella pneumophila* str. 2531STDY5467466 | FJEU01 | European Nucleotide Archive |
| *Legionella pneumophila* str. 2531STDY5467469 | FJEV01 | European Nucleotide Archive |
| *Legionella pneumophila* str. 2531STDY5467471 | FJEW01 | European Nucleotide Archive |
| *Legionella pneumophila* str. 2531STDY5467456 | FJEX01 | European Nucleotide Archive |
| *Legionella pneumophila* str. 2531STDY5467457 | FJEY01 | European Nucleotide Archive |
| *Legionella pneumophila* str. 2531STDY5467464 | FJEZ01 | European Nucleotide Archive |
| *Legionella pneumophila* str. 2531STDY5467465 | FJFA01 | European Nucleotide Archive |
| *Legionella pneumophila* str. 2532STDY5467639 | FJMD01 | European Nucleotide Archive |
| *Legionella pneumophila* str. ATCC 43283 | JFID01 | European Nucleotide Archive |
| *Legionella pneumophila* str. ATCC 33156 | JFIM01 | European Nucleotide Archive |
| *Legionella pneumophila* str. ATCC 43283 | LAXR01 | European Nucleotide Archive |
| *Legionella pneumophila* str. SH135 | LAYB01 | European Nucleotide Archive |
| *Legionella pneumophila* str. WX2011046 | LBAK01 | European Nucleotide Archive |
| *Legionella pneumophila* str. WD_4_1102b-36 | LBAN01 | European Nucleotide Archive |
| *Legionella pneumophila* str. ATCC 33156 | LBAV01 | European Nucleotide Archive |
| *Legionella pneumophila* str. NCTC12000 | UGOL01 | European Nucleotide Archive |
| *Legionella pneumophila* str. NCTC12024 | UGOM01 | European Nucleotide Archive |

**SUPPLEMENTARY TABLE 2**

The bacterial strains and plasmids used in this study.

| Name | Description |
| --- | --- |
| *Legionella pneumophila* str. Lens | The wild type strain of *L. pneumophila* str. Lens |
| SD-L1 | *Legionella pneumophila* str. Lens transformed with the SD-E3 plasmid |
| SD-L3 | *Legionella pneumophila* str. Lens transformed with the SD-E1 plasmid |
| SD-L5 | *Legionella pneumophila* str. Lens transformed with the SD-E5 plasmid |
| SD-L6 | *Legionella pneumophila* str. Lens transformed with the SD-E6 plasmid |
| SD-L8 | *Legionella pneumophila* str. Lens transformed with the SD-E12 plasmid |
| SD-L12 | *Legionella pneumophila* str. Lens transformed with the SD-E22 plasmid |
| SD-L56 | *Legionella pneumophila* str. Lens with the chromosomal CRISPR-Cas array deleted, save for the last repeat. Independent clone 1. |
| SD-L57 | *Legionella pneumophila* str. Lens with the chromosomal CRISPR-Cas array deleted, save for the last repeat. Independent clone 2. |
| SD-L58 | SD-L56 transformed with the SD-E5 plasmid |
| SD-L59 | SD-L57 transformed with the SD-E5 plasmid |
| SD-E1 | Priming plasmid with the protospacer sequence for the chromosomal Lens spacer 1 on the DNA (-) strand, pMMB207 vector backbone with chloramphenicol resistance |
| SD-E3 | Priming plasmid with the protospacer sequence for the plasmid Lens spacer 1 on the DNA (-) strand, pMMB207 vector backbone with chloramphenicol resistance |
| SD-E5 | Priming plasmid with the protospacer sequence for the plasmid Lens spacer 50 on the DNA (-) strand, pMMB207 vector backbone with chloramphenicol resistance |
| SD-E6 | Priming plasmid with the protospacer sequence for the chromosomal Lens spacer 23 on the DNA (-) strand, pMMB207 vector backbone with chloramphenicol resistance |
| SD-E12 | Priming plasmid with the protospacer sequence for the plasmid Lens spacer 50 on the DNA (+) strand, pMMB207 vector backbone with chloramphenicol resistance |
| SD-E22 | Priming plasmid with a scrambled protospacer sequence on the DNA (-) strand, pMMB207 vector backbone with chloramphenicol resistance |
| SD-E46 | pJB4648 vector backbone with an insert designed to knock out the chromosomal Lens CRISPR array, save for the last repeat, through allelic replacement. Gentamycin resistant with a sacB marker for sucrose selection. |

**SUPPLEMENTARY TABLE 3**

The oligos used in this study.

| Name | Sequence (5'-3') | Description |
|---|---|---|
| Lens-IF-pool-F | AGTTTGGTCAAATGCTTCAATA CTT | Used to screen *L. pneumophila* Lens samples for spacer acquisition, primer on leader sequence and has 100% identity for both chromosomal and plasmid Lens systems so can be used to screen for both systems. |
| pLens-IF-R | AGGATCATTAGAAAATCTTTTA CGC | Used to screen *L. pneumophila* Lens samples for spacer acquisition on the plasmid Lens I-F system, primer on 3rd spacer in CRISPR array. |
| cLens-IF-R | ATTCACATTTAACTCCAAATTT TGTT | Used to screen *L. pneumophila* Lens samples for on the chromosomal Lens I-F system, primer on 3rd spacer in CRISPR array. |
| pLens-IF-sp1(-)_F | CCCTTTTAAAAACCAATTTTTG CCGTGGCTCGATGGGCTGCA | The insert for the plasmid Lens spacer 1 targeted sequence on the DNA (-) strand. Contains PAMs on both side of the insert and ApaI/PstI overhangs when annealed with the complementary oligo. |
| pLens-IF-sp1(-)_R | GCCCATCGAGCCACGGCAAAA ATTGGTTTTTAAAAGGGGGCC | Complementary oligo to pLens-IF-sp1(-)_F. |
| pLens-IF-sp50(-)_F | CCCAATACGTGGTTATGACCTC ATCTAAAGACGTAGGCTGCA | The insert for the plasmid Lens spacer 50 targeted sequence on the DNA (-) strand. Contains PAMs on both side of the insert and ApaI/PstI overhangs when annealed with the complementary oligo. |
| pLens-IF-sp50(-)_R | GCCTACGTCTTTAGATGAGGTC ATAACCACGTATTGGGGGCC | Complementary oligo to pLens-IF-sp50(-)_F. |
| pLens-IF-sp50(+)_F | CCCTACGTCTTTAGATGAGGTC ATAACCACGTATTGGCTGCA | The insert for the plasmid Lens spacer 50 targeted sequence on the DNA (+) strand. Contains PAMs on both side of the insert and ApaI/PstI overhangs when annealed with the complementary oligo. |
| pLens-IF-sp50(+)_R | GCCAATACGTGGTTATGACCTC ATCTAAAGACGTAGGGGGCC | Complementary oligo to pLens-IF-sp50(+)_F. |
| cLens-IF-sp1(-)_F | CCCTTTTAAAAAACTTTAAGTT CTTTTCTGAAACAGGCTGCA | The insert for the chromosomal Lens spacer 1 targeted sequence on the DNA (-) strand. Contains PAMs on both side of the insert and ApaI/PstI overhangs when annealed with the complementary oligo. |
| cLens-IF-sp1(-)_R | GCCTGTTTCAGAAAAGAACTT AAAGTTTTTTAAAA GGGGGCC | Complementary oligo to cLens-IF-sp1(-)_F. |
| cLens-IF-sp23(-)_F | CCCTGATTGGTTTCTTTTTCTTG ACTGTTCTTTGAGGCTGCA | The insert for the chromosomal Lens spacer 23 targeted sequence on the DNA (-) strand. Contains PAMs on both side of the insert and ApaI/PstI overhangs when annealed with the complementary oligo. |
| cLens-IF-sp23(-)_R | GCCTCAAAGAACAGTCAAGAA AAAGAAACCAATCAGGGGGCC | Complementary oligo to cLens-IF-sp23(-)_F. |

| | | |
|---|---|---|
| Scrambled-IF-control_F | GCCTTATTAATCTAGAGTCGCC TTCTGAATCGAGTGGGGGCC | The insert for scrambled control plasmid. Contains PAMs on both side of the insert and ApaI/PstI overhangs when annealed with the complementary oligo. |
| Scrambled-IF-control_R | CCCACTCGATTCAGAAGGCGA CTCTAGATTAATAAGGCTGCA | Complementary oligo to Scrambled-IF-control_F. |
| pMMB207_pool_F | GATGCCCTCATTCAGCATTT | Used to screen the priming plasmid prior to use in transformations to ensure the insert was present after restriction enzyme cloning. Also used as the primer for Sanger sequencing to confirm the inserted sequence in the priming plasmid did not have mutations prior to the transformations. |
| pMMB207_pool_R | CACTTCTGAGTTCGGCATGG | Used to screen the priming plasmid prior to use in transformations to ensure the insert was present after restriction enzyme cloning. |
| cLens-array-KO-up-1kb-NotI-F | ATGGTCGACTTCTGGAGGTTGC TGGACATTATGCTCTGA | Used to design the insert to knock out the chromosome Lens CRISPR array. Has 1 Kb of homology upstream of the CRISPR array and a NotI site added for cloning into the pJB4648 vector. |
| cLens-array-KO-up-R | GGCAGTGAACTGTAATATAAA TCTTAAAAAAATAGTTACAAC AGAAAGTT | Complimentary oligo to the cLens-array-KO-up-1kb-NotI-F primer. Has homology to the cLens-array-KO-down-F primer added to it to facilitate the annealing of the upstream and downstream PCR products to create a single insert. |
| cLens-array-KO-down-2kb-SalI-R | ATGGCGGCCGCCGGTATCTTCT GCTGGACCA | Used to design the insert to knock out the chromosome Lens CRISPR array. Has 2 Kb of homology downstream of the CRISPR array and a SalI site added for cloning into the pJB4648 vector. |
| cLens-array-KO-down-F | TTATATTACAGTTCACTGCCGC ACAGGCAGCTTAGAAGTTATT ATGTTAA | Complimentary oligo to the cLens-array-KO-down-2kb-SalI-R primer. Has homology to the cLens-array-KO-up-R primer added to it to facilitate the annealing of the upstream and downstream PCR products to create a single insert. |
| cLens-screen-up-F | AACTGGCTGGTGGAGATATC | Used to screen the chromosome Lens array for whole array deletion following allelic replacement. |
| cLens-screen-down-R | CTGCCTGTTGAGTGACGTTG | Complementary oligo to cLens-screen-up-F. Also used with Lens-IF-pool-F to screen for spacer acquisition in the depleted array following cross-priming. |

**SUPPLEMENTARY TABLE 4**

The number of acquired spacers for the chromosomal Lens CRISPR-Cas system following array replenishment via cross-priming.

| Deletion Mutant | Total number of acquired spacers | % of new spacers that map to priming plasmid |
|---|---|---|
| Clone 1 | 738 | 98.92 |
| Clone 2 | 2726 | 98.86 |

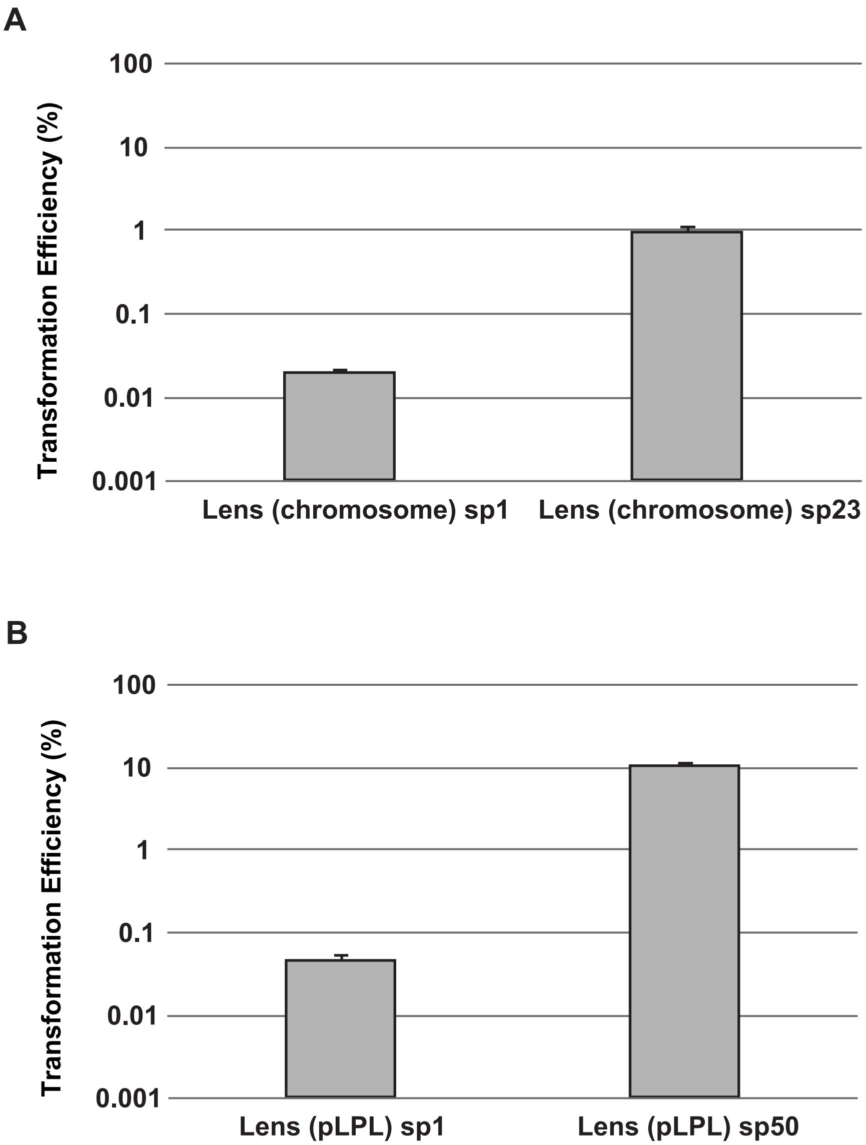


**Supplemental Figure 1 | The *L. pneumophila* str. Lens chromosome and pLPL type I-F CRISPR-Cas systems are active against plasmids containing protospacers.** *L. pneumophila* str. Lens was transformed with plasmids containing targeted protospacer sequences matched to the first spacer (sp1) or a mid-array spacer (sp23 or sp50, respectively) of the Lens (chromosome) CRISPR-Cas system **(A)** or the Lens (pLPL) CRISPR-Cas system **(B)**. After plating on selective media and incubating for three days, transformation efficiencies were calculated as a percentage of the transformation efficiency of a control plasmid with a scrambled targeted sequence. The average for three biological replicates is shown where the error bars represent the standard error of the mean.

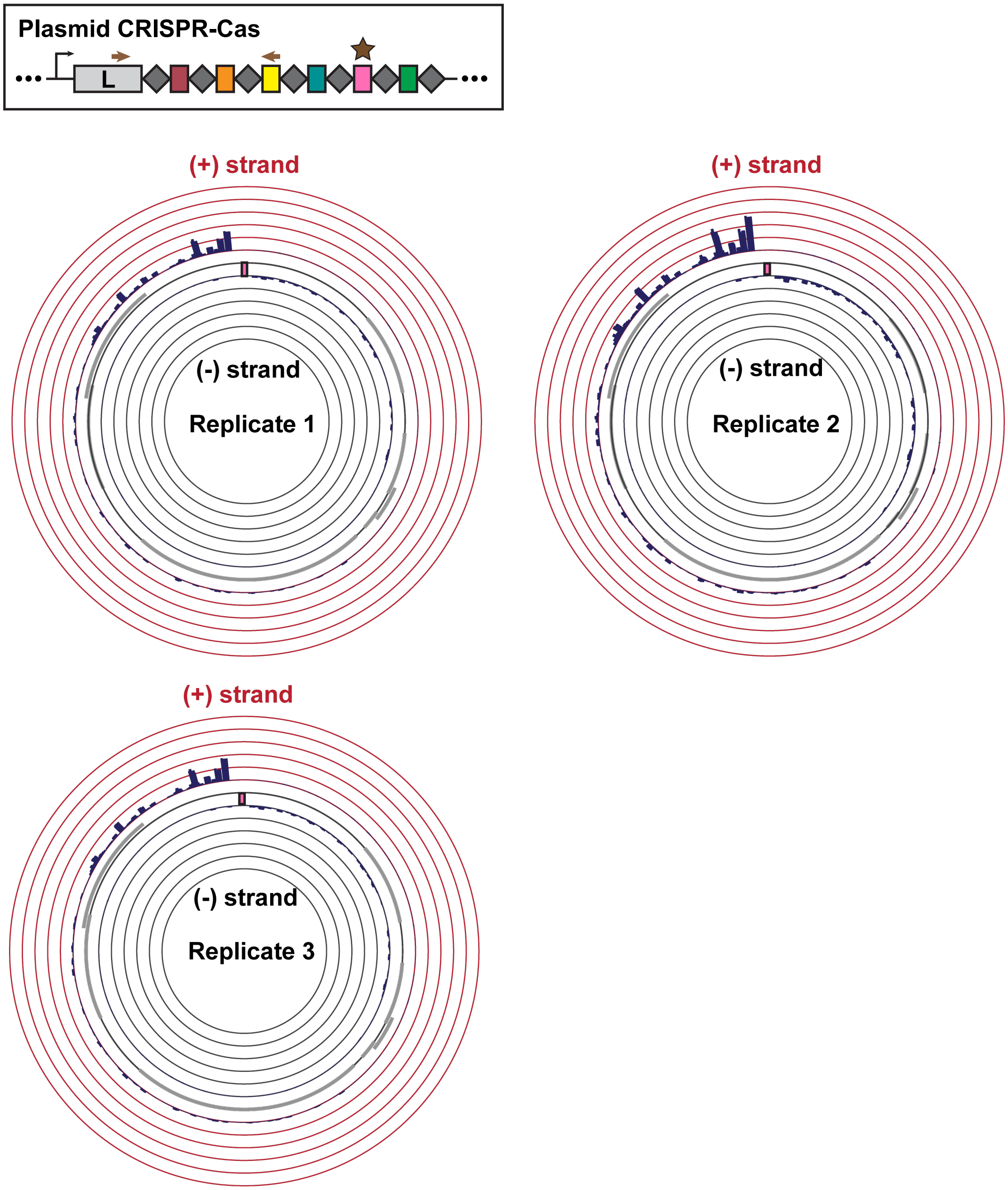


**Supplemental Figure 2 | Circos plots for the three replicates of targeted acquisition in the Lens (pLPL) CRISPR-Cas system.** Bacterial transformants with targeted plasmids were passaged for 20 generations without antibiotic selection to enrich for spacer acquisition; the leader-end of the CRISPR array was amplified and the amplicons were Illumina sequenced. Newly targeted protospacers were obtained from the raw reads using an in-house bioinformatics pipeline and were visualized with Circos. The height of the bars on the plot indicate the number of spacers mapped to the position on the plasmid, up to a 5% of total acquired spacers.

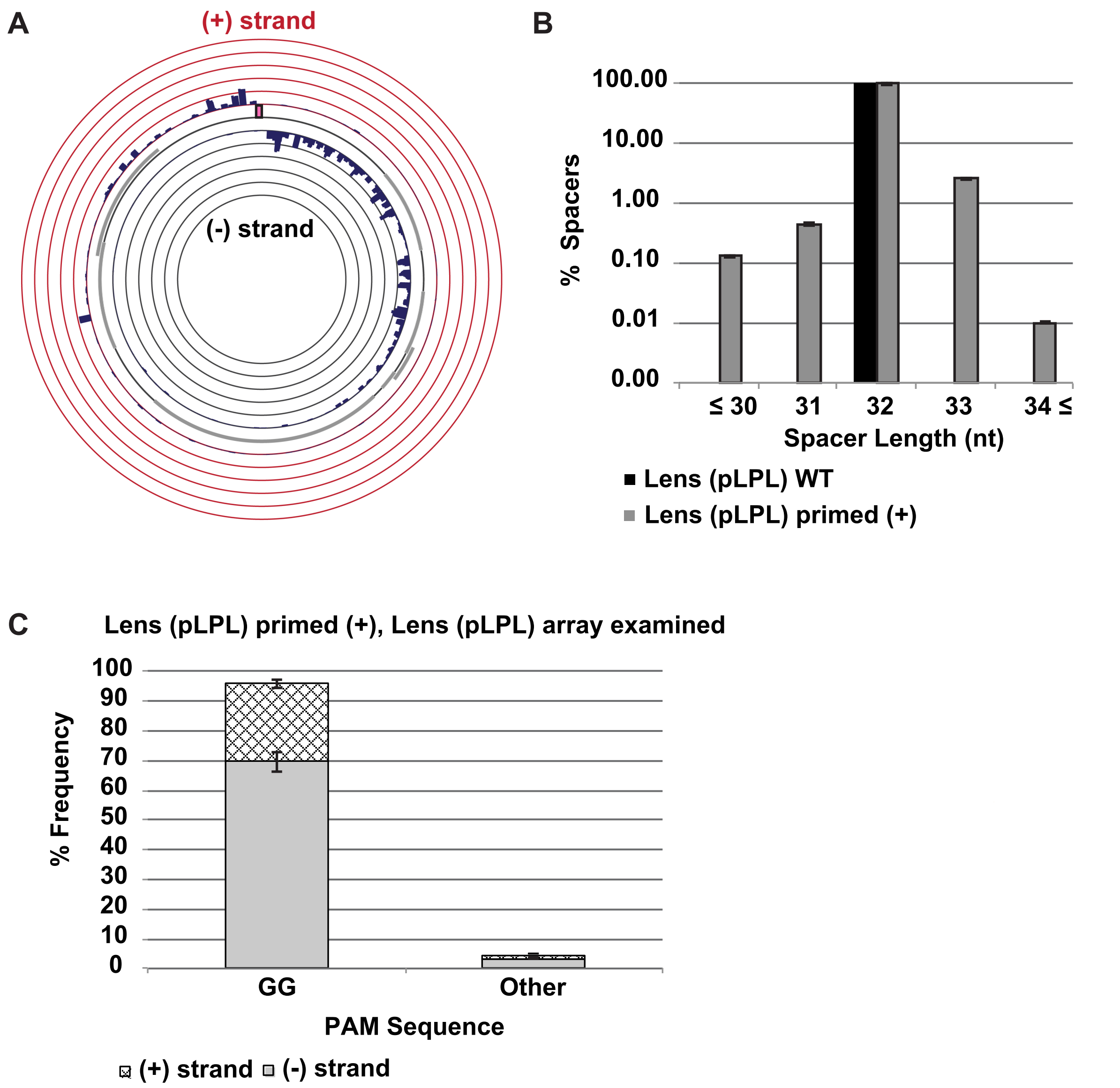


**Supplemental Figure 3 | The targeted spacer acquisition distribution for the Lens (pLPL) system when the priming protospacer is on the (+) strand.** The experimental set-up is the same as described for supplemental figure 2. **A)** Reversing the direction of the targeted sequence on the priming plasmid creates a mirrored distribution bias in acquired protospacers. **B)** The distribution of spacer lengths acquired from the targeted plasmid compared to the wild-type CRISPR-Cas array (n = 53). **C)** Quantification of the PAMs for the new protospacers in a stacked bar plot.

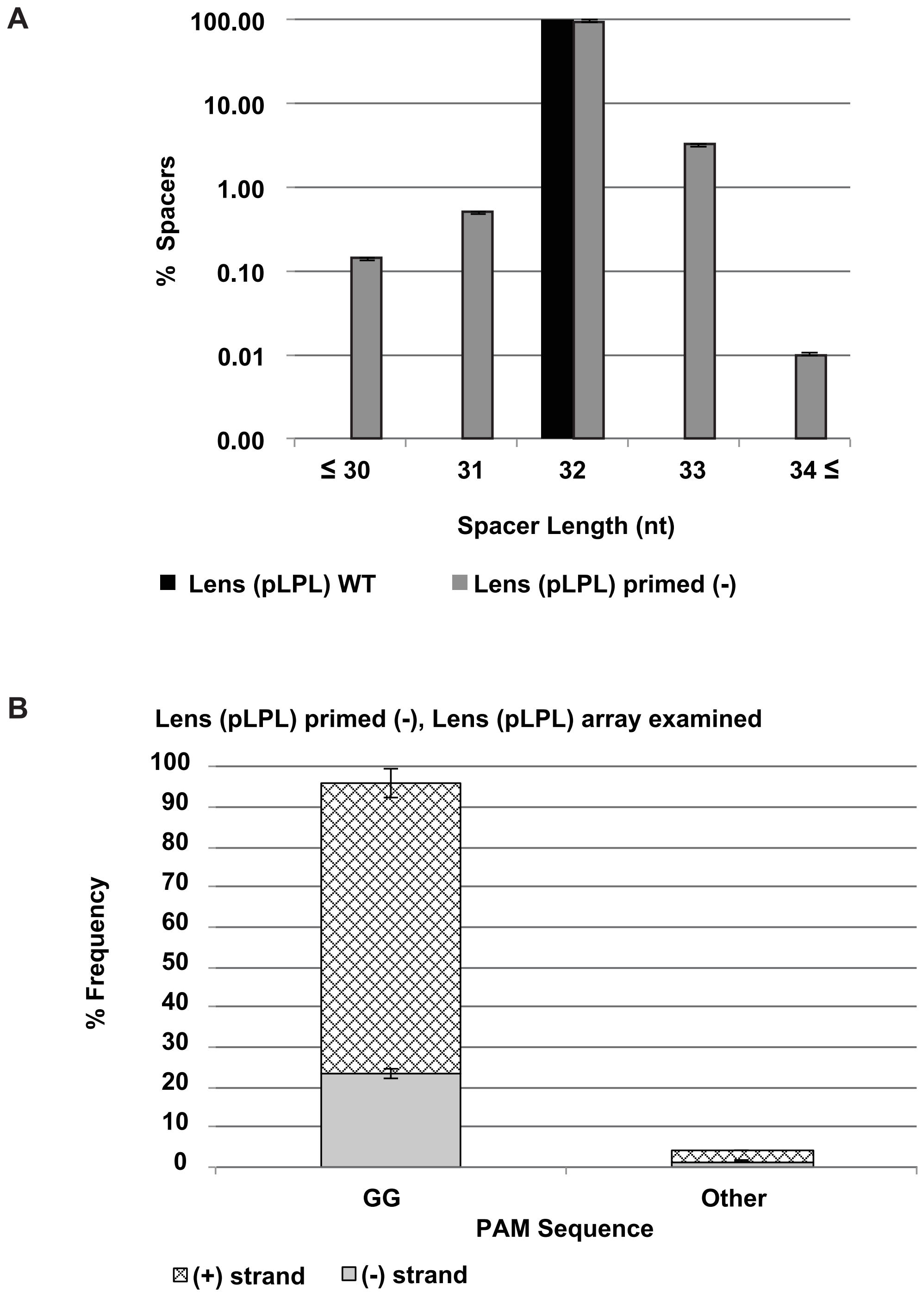


**Supplemental Figure 4 | Spacer length and PAM distributions for the Lens (pLPL) CRISPR-Cas system.** Experimental set-up is as described for supplemental figure 2. **A)** The distribution of spacer lengths acquired from the targeted plasmid (grey) compared to the wild-type CRISPR-Cas array (black, n = 53). **B)** Quantification of the PAMs for the new protospacers are shown in a stacked bar plot; the (+) strand PAMs are denoted by the hatched bars and the (-) strand PAMs are denoted by the grey bars.

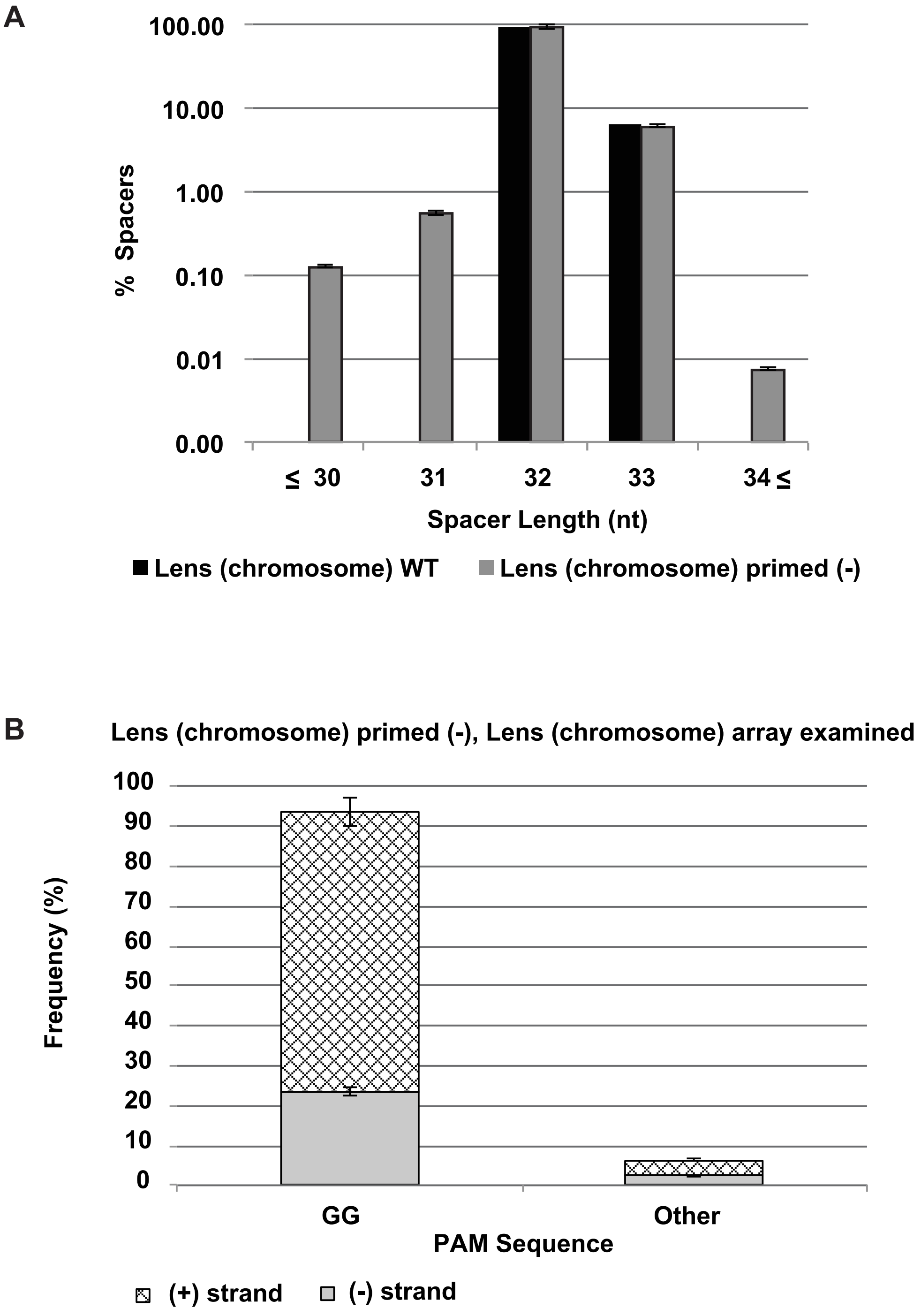


**Supplemental Figure 5 | Spacer length and PAM distributions for the Lens (chromosome) CRISPR-Cas system.** Experimental set-up is as described for supplemental figure 2. **A)** The distribution of spacer length acquired from the targeted plasmid compared to the wild-type CRISPR-Cas array (n = 64). **B)** Quantification of the PAMs for the new protospacers in a stacked bar plot.
